# The crustacean *Armadillidium vulgare* (Latreille, 1804) (Isopoda: Oniscoidea), a new promising model for the study of cellular senescence

**DOI:** 10.1101/583914

**Authors:** Charlotte Depeux, Ascel Samba-Louaka, Thomas Becking, Christine Braquart-Varnier, Jérôme Moreau, Jean-François Lemaître, Tiffany Laverre, Hélène Paulhac, François-Xavier Dechaume-Moncharmont, Jean-Michel Gaillard, Sophie Beltran-Bech

## Abstract

Senescence, the decline of physiological parameters with increasing age, is a quasi-ubiquitous phenomenon in the living world. The observed patterns of senescence, however, can markedly differ across species and populations, between sexes, and even among individuals. To identify the drivers of this variation in senescence, experimental approaches are essential and involve the development of tools and new study models. Current knowledge of the senescence process is mostly based on studies on vertebrates and main information about senescence in invertebrates is mostly limited to model organisms such as *Caenorhabditis elegans* or *Drosophila melanogaster*. In this context, we tested whether biomarkers of vertebrate ageing could be used to study senescence in a new invertebrate model: the common woodlouse *Armadillidium vulgare* (Latreille, 1804). More specifically, we looked for the effect of age in woodlouse on three well-established physiological biomarkers of ageing in vertebrates: immune cells (cell size, density, and viability), β-galactosidase activity, and the gene expression of telomerase reverse transcriptase (TERT), an essential subunit of telomerase protein. We found that the size of immune cells was higher in older individuals, whereas their density and viability decreased, and that the β-galactosidase activity increased with age, whereas the TERT gene expression decreased. These findings demonstrate that woodlouse displays age-related changes in biomarkers of vertebrate senescence, with different patterns depending on gender. The tools used in studies of vertebrate senescence can thus be successfully used in studies of senescence of invertebrates such as the woodlouse. The application of commonly used tools to new biological models offers a promising approach to assess the diversity of senescence patterns across the tree of life.

## INTRODUCTION

Many hypotheses have tried to explain why senescence is a quasi-ubiquitous phenomenon in the living organisms. For instance, the disposable soma theory proposed the senescence process as a result of damages accumulation over time (Kirkwood, 1977). These damages are strongly influenced by the environment, leading to trade-offs between the different functions (e.g., between reproduction and somatic maintenance) and shaping a high diversity of senescence patterns across species and populations, among individuals, and between sexes. One current challenge is to understand the selective forces and mechanisms driving this diversity of senescence patterns.

At the cellular level, senescence corresponds to the cellular deterioration leading to stop the cellular cycle (Campisi & d’Adda di Fagagna, 2007). As ageing is associated with cellular senescence (Herbig *et al.*, 2006; Wang *et al.*, 2009; Lawless *et al.*, 2010), many biomolecular parameters potentially inform about senescence and can therefore be valuable tools for studying this process (Bernardez de Jesus & Blasco, 2012). For example, the evolution of the integrity and efficiency of immune cells is particularly relevant to study cellular senescence because a diminution of the number of effective immune cells with increasing age takes place in both vertebrates (e.g., Cheynel *et al.*, 2017) and invertebrates (e.g., Park *et al.*, 2011). Another marker used to study cellular senescence is the enzymatic activity of the β-galactosidase. This enzyme is a hydrolase that transforms polysaccharides into monosaccharides. The lysosomal activity of this enzyme is increased when the cell enters in senescence (Dimri *et al.*, 1995; Itahana *et al.*, 2007). This phenomenon occurs in senescent cells of many organisms ranging from humans (Gary & Kindell, 2005) to honeybees (Hsieh & Hsu, 2011). Another protein linked to the cellular senescence process is the telomerase, a ribonucleoprotein complex composed by two essential components, the telomerase reverse transcriptase (TERT) and the telomerase RNA (TR) and other accessorial proteins (Podlevsky *et al.*, 2007). Telomerase lengthens the ends of telomeres (i.e., DNA sequences located at the end of chromosomes that protect chromosome integrity and shorten after each cell division). Cell senescence arises when the telomere length becomes critically short (Chiu & Harley, 1997; Shay & Wright, 2005). Telomerase activity depends on the species, age, as well as the type of tissue (e.g., Gomes *et al.*, 2010). For instance, telomerase is active during the development before birth and after only in stem and germ cells in humans (Liu *et al.*, 2007; Morgan, 2013) whereas in *Daphnia pulicaria* (Forbes, 1893) the telomerase activity in all tissues of the body decreases with increasing age (Schumpert *et al.*, 2015). The TERT is essential in the telomerase protein complex and has been shown to be related to cell survival in humans (Cao *et al.*, 2002). The TERT has also been detected in numerous species including vertebrates, fungi, ciliates, and insects (Robertson & Gordon, 2006; Podlevsky *et al.*, 2007).

Because the patterns of senescence are strongly diversified within the living world, it seems essential to study organisms displaying markedly different life history strategies to understand the causes and mechanisms underlying this diversity. Thus, invertebrates are increasingly used in experimental studies of senescence (Stanley, 2012; Ram & Costa, 2018). In addition to share similarities with vertebrates in terms of senescence, they can be manipulated experimentally and they are easier to monitor throughout their entire lifetime (Ram & Costa, 2018). These advantages make them models of choice for studying senescence. Here, we propose the common woodlouse *Armadillidium vulgare (*Latreille, 1804) as a promising new model for studying senescence. Woodlouse can live beyond three years and display sex-specific senescence patterns in natural populations (Paris & Pitelka, 1962). In addition, one study has already reported evidence of immune-senescence in this species (Sicard *et al.*, 2010).

In this context, we tested the suitability of β-galactosidase activity, immune cell parameters, and the TERT gene expression to cause age-specific responses in *A. vulgare.* According to reports in the literature, we expected an increase in β-galactosidase activity, and a decrease of both TERT gene expression and immune cell viability and density in *A. vulgare*. As males have higher adult survival than females (Paris & Pitelka, 1962), cellular senescence patterns are also expected to be sex-specific in this species.

## MATERIALS AND METHODS

### Biological model

Individuals of *A. vulgare* used in the experiments were obtained from a wild population collected in Denmark in 1982. These individuals have been maintained on moistened soil under the natural photoperiod of Poitiers, France (46.58°N, 0.34°E) at 20 °C, and fed *ad libitum* with dried linden leaves and carrots. Crosses were monitored to control and promote genetic diversity. For each clutch obtained, individuals were sexed, and brothers and sisters were separated to ensure virginity. Woodlice are promiscuous and only breed when they are one-year-old during their first breeding season in the spring. Females carry their offspring in a ventral pouch (marsupium) and can produce up to three clutches per season. In the common woodlouse, individuals molt throughout their lifetimes, with approximate one molt per month. During this process all the cells of the concerned tissues are renewed (Steel, 1980). The brain, the nerve cord, and gonads, however, are not renewed during molting and are therefore relevant candidates for tissue-specific study of senescence in this species. Woodlice were classified in three different age categories: juvenile (before first reproduction, birth to one-year-old), young (one-year-old, first year of reproduction), and old (from and over two-year-old), which are very rare in natural populations (Paris & Pitelka, 1962). As the woodlouse is continuous grower, juveniles and small individuals could not be used for certain experiments, especially when they required protein extraction or hemolymph collection. Thus, we were only able to test in them the telomerase expression. Old individuals were sampled according to the number of individuals available in our breeding; sometimes we had two-year-old individuals and sometimes three-year-old ones. Moreover, males and females were tested separately to assess the impact of sex.

### Measurement of β-galactosidase activity

To test the impact of age on β-galactosidase activity, 180 individuals were used: 90 young (i.e., six-month old, 45 males, and 45 females) and 90 old (two-year-old, 45 males, and 45 females).

Individuals were dissected separately in Ringer solution (sodium chloride (NaCl) 394 mM, potassium chloride (KCl) 2 mM, calcium chloride (CaCl_2_) 2 mM, sodium bicarbonate (NaHCO_3_) 2 mM) and nerve cord was removed. To obtain a sufficient amount of protein, we made pools of five nerve cords from five different individuals of the same age. The five nerve cords were filed in 500 μl of Lyse buffer 1X (Chaps detergent ((3-((3-cholamidopropyl) dimethylammonio)-1-propanesulfonate) 5 mM, citric acid (C_6_H_8_0_7_) 40 mM, sodium phosphate (Na_3_PO_4_) 40 mM, benzamidine (C_7_H_8_N_2_) 0.5 mM, and PMSF (phenymethylsulfonyle (C_7_H_7_FO_2_S) 0.25 mM, pH 6) (Gary & Kindell, 2005), and then centrifuged at 15,000g at 4 °C for 30 min. The supernatant was taken and kept at –80°C until its utilization. The protein concentration was determined by the BCA (bicinchoninic acid assay) assay (Thermo Fisher Scientific, Waltham, MA, USA) and homogenized at 0.1 mg ml ^−1^. The β-galactosidase activity was measured as described by Gary & Kindell (2005). Briefly, 100 μl of extracted protein at the concentration of 0.1 mg ml ^−1^ were added to 100 μl of reactive 4-methylumbelliferyl-D-galactopyranoside (MUG) solution in a 96-wellmicroplate. The MUG reactive, in contact to β-galactosidase, leads by hydrolysis to the synthesis of 4-methylumbelliferone (4-MU), which is detectable using fluorescent measurements. Measures were performed by the multimode Mithrax microplate reader (LB940 HTS III, excitation filter: 120 nm, emission filter 460 nm; Berthold Technologies, Bad Wildbad, Germany) for 120 min. Two technical replicates were measured for each nerve pool.

### Measurement of immune cell parameters

To test the impact of age on the immune cell parameters (i.e., density, viability, and size), we were able to undertake individual tests because of our previous experience. It was therefore not necessary to carry out pools of hemolymph unlike proteins or DNA to make the measurements. Sixty mature individuals were used: 30 young (i.e., one-year-old, 15 males, and 15 females) and 30 old (three-year-old, 15 males, and 15 females) individuals.

To study the impact of age on the immune parameters, a hole was bored in the middle of the sixth segment and 3 μl of haemolymph were collected per individual with an eyedropper and deposited promptly in 15 μl of anticoagulant solution (MAS-EDTA (EDTA (ethylenediaminetetraacetic (C_10_H_16_N_2_0_8_)) 9 mM, trisodium citrate (Na_3_C_6_H_5_O_7_) 27 mM, sodium chloride (NaCl) 336 mM, glucose (C_6_H_12_O_6_) 115 mM, pH 7 (Rodriguez *et al.*, 1995)). Then, 6 μl of trypan blue at 0.4% (Invitrogen, Carlsbad, CA, USA) were added to color the dead cells. Thereafter, 10 μl of this solution were deposed in a counting slide (Invitrogen Coutness®; Thermo Fisher Scientific). The immune cell density, the immune cell viability, and the immune cell size were evaluated using an automated cell counter (Invitrogen Countess®).

### Measurement of TERT gene expression

The identification of the telomerase reverse transcriptase (TERT) gene was first performed from the *A. vulgare* genome (Chebbi *et al.*, 2019). In order to test whether this gene was present and preserved in crustaceans, phylogenetic analyses were undertaken upstream (see Supplementary material S1–S4). This gene has been found in crustacean transcriptomes and the topology of the TERT-gene tree follows the phylogenetic relationships between the involved species (Supplementary S3), suggesting a conserved role of the TERT gene.

### Gene expression

We tested the effect of age on the expression of TERT gene within four different age groups: 1) four-month old, 2) one-year-old, 3) two-year-old, and 4) three-year-old. Females and males were tested separately by pools of five individuals in one-, two-, and three-year-old groups and by pools of seven individuals in a four-month-old group as smaller individuals provide less biological material. All conditions required four replicates for each sex. A total of 176 individuals were used for this experiment. For each group we tested the expression level of the TERT gene in two different tissues: the nerve cord (somatic line) and gonads (germinal line).

Animals were washed by immersion for 30 seconds in a 30% sodium hypochlorite solution (NaClO) followed by two 30-sec immersions in distilled water. Tissues were dissected in Ringer solution (sodium chloride 394 mM, potassium chloride (KCl) 2 mM, calcium chloride (CaCl_2_) 2 mM, sodium bicarbonate (NaHCO_3_) 2 mM) and deposited by pools of five units for each tissues on TRIzol reagent (Invitrogen) to extract RNA according to the manufacturer’s protocol after a cell disintegration using a Vibra Cell 75,185 sonicator (amplitude of 30%). Total RNA was quantified by NanoDrop technology and stored at –80 °C until use. Reverse transcriptions (RT) were made from 500 ng of RNA previously extracted and using the SuperScript^TM^ IV Reverse Transcriptase kit (Thermo Fisher Scientific) according to the manufacturer’s instructions. Primers were designed using the identified gene: primer TERT_F: 5’-AGGGAAAACGATGCACAACC-3’ and primer TERT_R: 5’-GTTCGCCAAATGTTCGCAAC-3’ (see Supplementary material S1). Quantitative RT-PCR was performed using 0.6 μl of each primer (10 μM), 2.4 μl of nuclease-free water, and 1.5 μl of cDNA template and the LightCycler LC480 system (Roche, Pleasanton, CA, USA) as follows: 10 min at 95 °C, 45 cycles of 10 s at 95 °C, 10 s at 60 °C, and 20 s at 72 °C. Expression levels of target genes were normalized based on the expression level of two reference genes previously established: the ribosomal protein L8 (RbL8) and the Elongation Factor 2 (EF2) (Chevalier *et al.*, 2011).

### Statistical analyses

All statistical analyses were performed using the R software v.3.5.2 (R. Core Team, 2018). The β-galactosidase activity was analyzed with linear mixed effect models using the package lme4 (Bates *et al.*, 2014). As two technical replicates were measured for each pool, the model included the pools fitted as a random effect, age and sex and their two-way interaction as fixed factors.

Concerning the immune parameters, linear models with Gaussian distribution were fitted to analyze variation in the cell size and viability. A linear model of the cell number (log-transformed; (Ives, 2015) was fitted for the cell density.

The level of TERT expression according to age in the two different tissues were compared by a Kruskal-Wallis rank sum test in combination with Nemenyi’s post hoc multiple comparison test with the Tuckey correction using R package PMCMR (Pohlert, 2014).

## Results

### β-galactosidase activity

The β-galactosidase activity was higher in old (i.e., two-year-old) than in young (i.e., six-month old) individuals (χ^2^_1_ = 6.15, *P* = 0.013; Fig. 1). We also detected a higher β-galactosidase activity in females than in males (χ^2^_1_ = 7.26, *P* = 0.007; Fig. 1).

**Figure 1.**
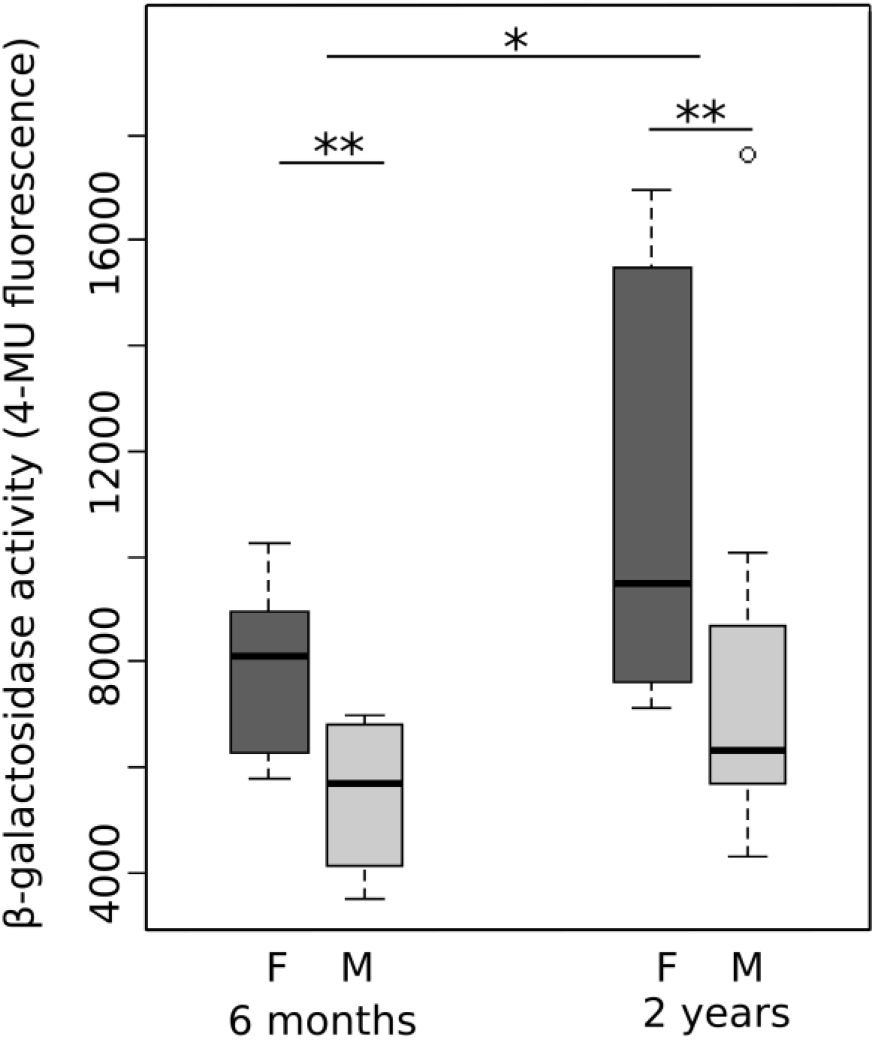
Boxplot of β-galactosidase activity in *Armadillum vulgare* according to age and sex (F, females, M, males). The thick line represents the median, the box the interquartile range, and the whisker the most extreme data point within 1.5 of the interquartile range. The outliers outside this range are shown as open circles. N = 24 pools of 5 individuals; * *P* < 0.05, ** P<0.01.

**Figure 1.**
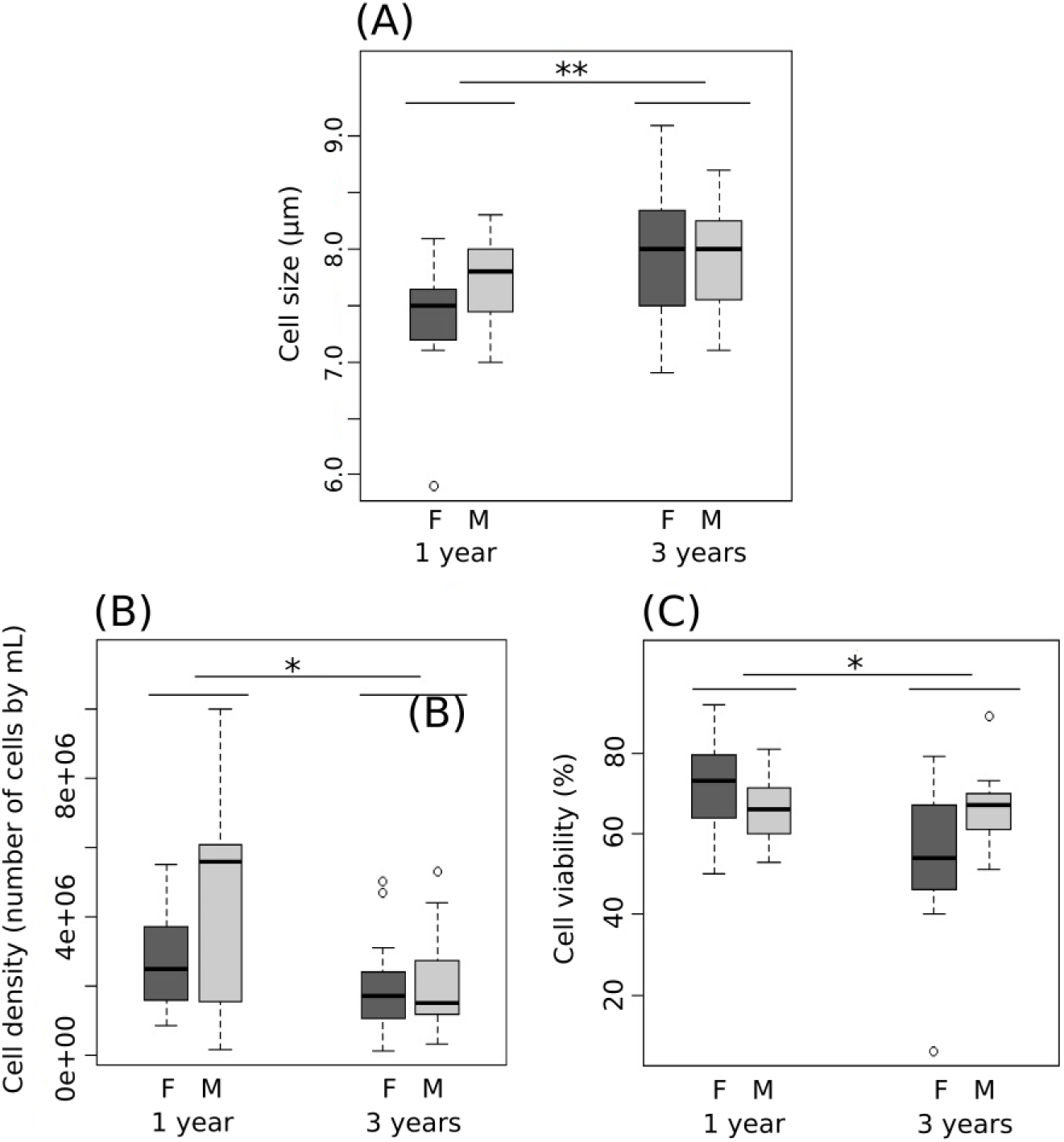
Immune cell size (**A**), density (**B**), and viability (**C**) according to age and sex in *Armadillum vulgare* (F, females; M, males). The thick line represents the median, the box the interquartile range, and the whisker the most extreme data point within 1.5 of the interquartile range. The outliers outside this range are shown as open circles. N = 60 individuals: 15 one-year-old females, 15 one-year-old males, 15 three-years-old females, and 15 three-years-old males; * *P* < 0.05, ** *P* < 0.01.

### Immune cells parameters

Cell size was higher in three-year-old than in one-year-old individuals (F_1,58_ = 8.54, *P* = 0.005; Fig. 2A). Conversely, the cell density was higher in one-year-old than in three-year-old individuals (F_1,58_ = 4.33, *P* = 0.01; Fig. 2B). Concerning the immune cell viability, a statistically significant interaction occurred between age and sex, with a relatively lower immune cell viability in three-year-old females (F_3,56_ = 6.85, *P* = 0.01; Fig. 2C). No sex effect was detected on cell size (F_2,57_ = 0.76, *P* = 0.38; Fig. 2A) or cell density (F_2,57_ = 0.32, *P* = 0.57, Fig. 2B).

**Figure 2.**
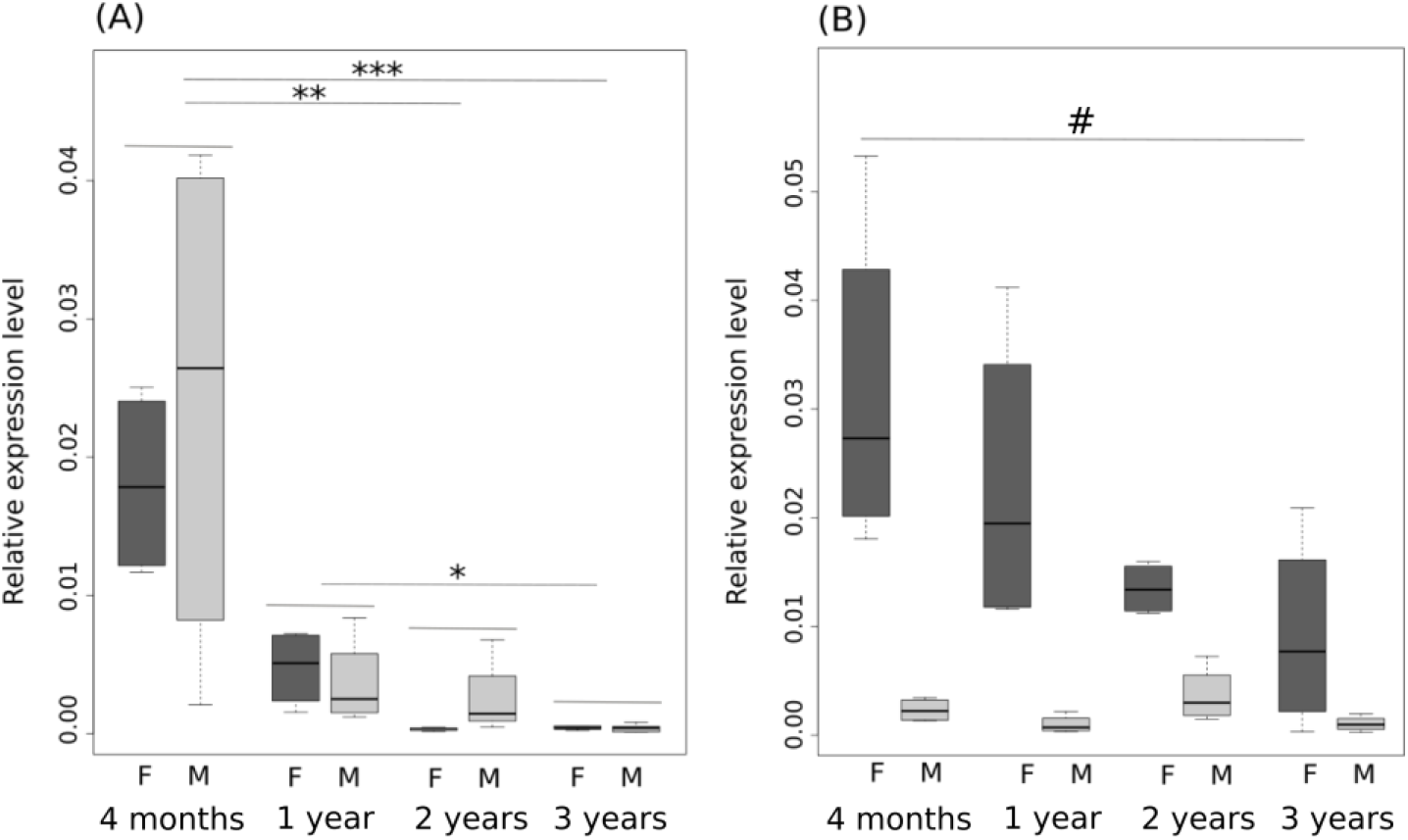
Relative expression level of TERT expression in nerve cords (A) and in gonads (B) in *Armadillum vulgare* (F, females; M, males). Expression of each gene was normalized based on the expression of ribosomal protein L8 (RbL8) and elongation factor 2 (EF2) as reference genes. The thick line represents the median, the box the interquartile range, and the whisker the most extreme data point within 1.5 the interquartile range. N = 176 individuals: 28 four-month-old females, 28 four-month-old males, 20 one-year-old females, 20 one-year-old males, 20 two-year-old females, 20 two-year-old males, 20 three-year-old females, 20 three-year-old males; # *P* < 0.10, ** *P* < 0.01

### TERT gene expression

The TERT gene expression decreased with increasing age in nerve cords (χ^2^_3_ = 23.30, *P* < 0.001; Fig. 3A). TERT expression was higher in four-month-old individuals compared to two-year-old and three-year-old individuals (*P* = 0.001 in both cases) and in one-year-old individuals compared to three-years-old individuals (*P* = 0.038), without any detectable sex effect (χ^2^_1_ = 0.14, *P* = 0.70; Fig. 3A). In gonads, the TERT gene expression was much higher in females (χ^2^_1_ = 17.81, *P* < 0.001; Fig. 3B) and tended to decrease with increasing age (χ^2^_3_ = 7.5, *P* = 0.057; Fig., 3B) as the TERT gene expression tended to be higher in four-month-old females compared to three-year-old females (*P* = 0.054). A general tendency was also observed in males (χ^2^_1_ = 7.34, *P* = 0.061; Fig. 3B), the TERT gene expression tending to be higher in two-year-old individuals compared to one-year-old and three-year-old individuals (*P* = 0.14 and *P* = 0.12, respectively; Fig. 3B).

## Discussion

We tested several effective physiological biomarkers of vertebrate senescence to assess whether they could also be used in invertebrates such as the common woodlouse. Immune cells showed an increase in their size and a decrease in their density and viability with increasing age. The activity of the β-galactosidase enzyme in nerve cords increased. The TERT gene expression decreased with increasing age in nerve cords of males and females and in the gonads of females. In contrast, the TERT gene expression was very low in the male gonads to suggest a role on the cellular senescence status in this tissue. The difference regarding the expression of TERT in the nerve cords and gonads underlies the importance of organ choice to perform such an analysis.

Our study is in line with previous studies that had previously revealed the possibility of using vertebrate biomarkers in invertebrates (Hsieh & Hsu, 2011; Park *et al.*, 2011; Schumpert *et al.*, 2015). By testing a set of different physiological biomarkers of vertebrate senescence, often studied independently, our study supports both ideas that routinely used biomarkers in vertebrates can be adapted in invertebrates and that the senescence process is quasi-ubiquitous in the living world and can be expressed in a similar way in very different organisms.

Previous studies have shown that the probabilities to survive decrease with increasing age in *A. vulgare* (Paris & Pitelka, 1962). The cellular damages accumulated during the life of the isopod could be the cause of cell senescence and therefore the driving force behind actuarial senescence (Harman, 1956; Barja, 2000; Barja & Herrero, 2000; Finkel & Holbrook, 2000). In *A. vulgare*, the 2- and 3-year-old individuals could have therefore accumulated more cellular damages during their lifetime, leading to the cellular senescence we report.

Our study also revealed a strong difference between sexes on the response of biomarkers to age changes. At a given age, females display higher β-galactosidase activity and lower immune cell viability than males, as if they age faster than males. Between-sex differences in lifespan have been reported in *A. vulgare* with a longer lifespan in males than in females (Geiser, 1934; Paris & Pitelka, 1962). Exact differences in actuarial senescence patterns (i.e., age-specific changes in survival probabilities) remain to be quantified in *A. vulgare* but such differences are quite common both in vertebrates and invertebrates (Tidière *et al.*, 2015; Marais *et al.*, 2018). One of the main hypotheses proposed to explain sex differences in longevity or senescence patterns relies on different resource allocation strategies between sexes (Vinogradov, 1998; Bonduriansky *et al.*, 2008). Females carry their offspring in their marsupium for one month, giving nutrients and protection, thus, they allocate more energy to reproduction than males that do not provide any parental care. The fact that the lifespan of *A. vulgare* is shorter in females (Paris & Pitelka, 1962) thus supports a role of differential sex allocation in this species. Moreover, the difference between males in TERT expression in gonads also suggest a difference in sex allocation according both to age and sex. The increased in TERT expression in male gonad in two-years-old individuals should be the result of a terminal investment in reproduction for males: by increasing their TERT expression and thus potentially their telomerase activity they should provide a better fitness to their offspring. This allocation may be lost when males are too old (three-year-old).

Sex differences in resource allocation strategies could also be driven by environmental conditions (Shertzer & Ellner, 2002). Our physiological biomarkers of vertebrate senescence revealed sex differences, and as supported in Depeux *et al.* (2019), they could constitute useful tools to identify other factors involved in variations in senescence patterns, such as environmental stressors. Moreover, if these biomarkers seem to predict better the physiological age than chronological age, notably in terms of survival and reproduction, they could correspond to biomarkers of senescence in the woodlouse (Baker & Sprott, 1988; Simm *et al.*, 2008; Sprott, 2010).

The physiological biomarkers of vertebrate senescence thus respond to age changes in *A. vulgare*, a species that represents a new invertebrate model of ageing. The parameters that predict the chronological age of *A. vulgare* individuals might offer reliable biomarkers, especially if their measurements are related to both reproductive and survival prospects more than to the chronological age of individuals. The availability of its genome, the ease of its breeding, its particularity to continue growing during its lifespan, and its adaptations to terrestrial life as well as the presence of cellular senescence on our senescence biomarkers make it a promising candidate to study senescence.

## SUPPLEMENTARY MATERIAL

S1. Phylogenetic analysis protocol of the TERT gene in crustaceans.

S2. TERT-gene alignment used for the phylogenetic analysis.

S3. TERT-gene tree following the phylogenetic relationships between the species involved.

S4. TERT-gene tree following the phylogenetic relationships between the species involved (Newick format).

## Acknowledgements

We would like to thank Sylvine Durand, Richard Cordaux, Isabelle Giraud, and Bouziane Moumen for our constructive discussions as well as Marius Bredon, Carine Delaunay, Maryline Raimond, and Alexandra Lafitte for technical assistance. The preprint of this work has been deposited in BioRxiv (https://doi.org/10.1101/583914). This work was supported by the 2015– 2020 State-Region Planning Contract and European Regional Development Fund and intramural funds from the Centre National de la Recherche Scientifique (CNRS) and the University of Poitiers. JFL and JMG were supported by a grant from the Agence Nationale de la Recherche (ANR-15-CE32-0002-01 to JFL). A.S.L. is supported by a grant from the Agence Nationale de la Recherche (ANR-17-CE13-00001-01 “Amocyst”). Our research also received funding from the *Appel à projets de recherche collaborative inter-équipes 2016-2017* by the laboratory Ecologie et Biologie des Interactions.

