## Supplementary material S1 for "The crustacean *Armadillidium vulgare* (Latreille, 1804) (Isopoda: Oniscoidea), a new promising model for the study of cellular senescence"

### Supplementary material 1

**Phylogenetic analysis of the TERT gene in crustaceans**

### Gene identification and dataset preparation

Homologous sequences identification of TERT was then performed on the complete translated transcriptome datasets for oniscidean species previously generated by Becking *et al.* (2017) using BLASTp (version 2.7.1, Camacho *et al.*, 2009). Then, the search was extended to all the crustacean transcriptomes available in the Transcriptome Shotgun Assembly database (TSA, https://www.ncbi.nlm.nih.gov/genbank/tsa/) with tBLASTn (2.7.1, Camacho *et al.*, 2009), using protein sequences indentified in *A. vulgare* as query. A scan for the presence of Pfam domains (Bateman et al., 2004) using HMMER (http://hmmer.org/) (Finn et al., 2011) was performed, and only the amino acids sequences containing the Pfam Reverse-transcriptase domain (Pfam: PF00078) were retained. Finally, only the amino acid sequences having a total length ≥ 100 bp were considered.

### Phylogenetics analysis

A multiple alignment was performed with the Geneious aligner (version 7.0.6, Kearse *et al.*, 2012) using default parameters. These multiple alignment was then trimmed with GBLOCKS (version 0.91b, Talavera & Castresana, 2007) to remove ambiguously aligned regions, and to maximize the resulting alignment length, we used the less stringent options to obtain a 120 amino acids alignment length. The alignment fasta file is available in Supplementary Materials 2. To determine the best-fit model of nucleotide substitution, we used Prottest (version 3.4, Darriba*et al.*, 2011). The best substitution model was the LG+G model according to all information criteria. Maximum-likelihood analyses were performed independently on each protein alignment using RAxML (version 7.4.6,Stamatakis, 2006)with 100 independent replicates followed by 1000 replicates of bootstrap resampling. Bootstrap values were subsequently mapped onto the optimal consensus tree obtained from the 100 independent searches. Only nodes having a bootstrap value ≥ 50 were considered. The final tree files (newick format) is available in Supplementary Material 4.
