## Supplementary material S2 for "The crustacean *Armadillidium vulgare* (Latreille, 1804) (Isopoda: Oniscoidea), a new promising model for the study of cellular senescence"

### Supplementary material 2

>NP_001035796.1

---------- ---------- ----IYKWNP LSGCLHRTVD DYFFCSPHPH KVYDF-ELLI

KGVYQ----V NPTKTRTNLE IPYCGKIFNL TTRQPARFLQ KAMDFPFICN TVFNDQRTVF

>gb|GEHY01011681.1|:C2384-2022

SYLKLAKTAT YKVVQGIPQG WPLSETLCNL YYGEMFRGVD DFLYITTDHG KALRFLQTMC

QGIEEYGCSV ESKKVCTNL- ---------- ---------- ---------- ----------

>gb|GFUD01030199.1|:149-496

SYKLGSVKKT FLVTQGVIQG DPLSSNLSDI YYGDMLRGAD DFLFVSTSRR RVQSFHQFLT

SGLASHNCHF KPSKTKTNV- ---------- ---------- ---------- ----------

>gb|GEFB01013694.1|:C752-144

QPCVQYRNLK VKIAEGIPEG CPLSTELCNF FYASMIRTVD DFLYISRSLE NTQQFHDRMM

KGIHEFNCYI NEDKTLTNIR FYFNGYIICL VSECLHQTIG CCLPYQLIAP NVWRTGIICG

>gb|GGFD01023608.1|:1617-2150

---------- --IAQGLPQG GCLSTDLCNF YYARMIRIAD DFLFLSKSLE KAQEFQFRIK

SGIQEFNCQF NPKKTESNIR VYFNGYIICM FTKSLFQILN SRLKGPMMNH NVWRAGIHCG

>gi|936323717|gb|GDRN01098857.1|:1558-2268

---------- ---------- ------LCDF YLARMIRVAD DFLFLSKSLE KAQKFKFRMT

AGIADFNCFI NSKKTLSNIK IYFNGYIICM ITNTLFIIVR NRLQKHMMVP NVWRVGIHCG

>gi|936323711|gb|GDRN01098861.1|:1938-2648

---------- ---------- ------LCDF YLARMIRVAD DFLFLSKSLE KAQKFKFRMT

AGIADFNCFI NSKKTLSNIK IYFNGYIICM ITNTLFIIVR NRLQKHMMVP NVWRVGIHCG

>gi|936323715|gb|GDRN01098858.1|:2019-2729

---------- ---------- ------LCDF YLARMIRVAD DFLFLSKSLE KAQKFKFRMT

AGIADFNCFI NSKKTLSNIK IYFNGYIICM ITNTLFIIVR NRLQKHMMVP NVWRVGIHCG

>gb|GFFJ01110714.1|:1-405

---------- ---------- ---------- ---------- -FLFLSKSLE KAQEFRFRMM

KGIKEFNCHI NLKKTDSNIR IHFNGYIICM STKSLFIISK NRLQRHMIDP NVWRVGIHCG

>gi|595481066|gb|GARW01005962.1|:C1387-1037

MLKYGYQKLF YIVRKGIVVG DSLSSPLCDV YYGQLLRGMD DFIFVSTNQK EAERFLTVME

NGFPDYGCEI QKQKTKTNF- ---------- ---------- ---------- ----------

>gb|GEBG01023904.1|:C1377-466

QFFREGKNY- FSLICGLAQG GKLSFHLSDL YYSTMVKKAD DILFVSVLHK EAEKFLGRIQ

SGIPEFNCII NEAKTSHNVR TVFNGVVVCL HSRQLRQVSA ARLTLEVLNP NVWRTGVMAG

>gb|GFDA01078771.1|:C429-103

QFFREGKNY- FSLICGLAQG GKLSFHLSDL YYSTMVKKAD DILFVSVLHK EAEKFLGRIQ

SGIPEFNCII NEAKTSHNV- ---------- ---------- ---------- ----------

>gb|GFUC01026601.1|:1433-1759

QFFREGKNY- FSLICGLAQG GKLSFHLSDL YYSTMVKKAD DILFVSVLHK EAEKFLGRIQ

SGIPEFNCII NEAKTSHNV- ---------- ---------- ---------- ----------

>gb|GFFH01034216.1|:215-1102

QLVRNGQSQ- YLVTQGISQG GALSSLLCEL YYSAMLRSVD DFLYVSPHKD LAESFLQTMK

DGIMEYNCSI NELKTQTNLQ VTFTGSILCS DTCELLQVSV AKMAPLYIDA NVWRNSLLAA

>gb|GETT01116158.1|:C1474-671

QLIQCGPRQN FILIRGISQG GTLSSALCEL YYSSMLRSVD DMLFVTPDLK KARNFLQKME

AGFPDFNCKM NNFKTAHNIR VSFCGVVVCS TSHQLRQVTL ARLNTTVLDP NIWRTGVMAG

>lcl|IABX01122151.1

QIVHAGQKVR LLIRKGIPQG GRLSAALCNL YYSAFLRSVD DFLYVSASYD SASRYFDFMK

NGIPEFNSYI NEQKTFHNL- ---------- ---------- ---------- ----------

>gi|1066151538|emb|HACK01015471.1|:1814-2719

LIISNGSKL- FRIVRGLPQG GSLSAALCEF YYAAMFRAVD DFLFISEARR DAEQFMEHVK

SGIPEFNCSM NKEKTLHNLR VSFCGIIVCL KSLQLRQVTL TRLQPLMLDP NVWRSGFMAG

>gi|1068206298|emb|HACB02024865.1|:C917-87

---------- FRIVRGLPQG GSLSAALCEF YYAAMFRAVD DFLFISEARR DAEQFMEHVK

SGIPEFNCSM NKEKTLHNLR VSFCGIIVCL KSLQLRQVTL TRLQPLMLDP NVWRSGFMAG

>gb|GBEI01054168.1|:C570-235

QVITNGSLL- LRVACGVPQG GTLSAALCEF YYASMLRAVD DFLFISQTRD EAKRFLEQIV

CGVPDFNCHM NNEKTFHNL- ---------- ---------- ---------- ----------

>gi|703876552|gb|GBEV01020382.1|:C1209-304

QVITNGSLL- LRVARGVPQG GTLSAALCEF YYASMLRAVD DFLFISQSRD EAKRVLEQTV

CGVPDFNCHM NREKTFHNIR VPFCGVIVCL KSLQLQQVTL ARLKPLVLDP NVWRSGIMAG

>gi|742973106|gb|GARH01006483.1|:C786-1

---------- -----GVPQG GTLSAALCEF YYASMLRAVD DFLFISQSRD EAKRVLEQTV

CGVPDFNCHM NREKTFHNIR VPFCGVIVCL KSLQLQQVTL ARLKPLVLDP NVWRSGIMAG

>gb|GFRT01019308.1|:C999-88

QIAKVTQRQC IRANRGVPQG GSLSGALCEL YYSAMLRSVD DFLYVTPSSS QASLFLTRMR

KGFPDFNCFI NERKTQHNLR VVFRGIVMCS ESRQLQQVSL ARLPQVVLDP NVWRAGFMAG

>gb|GEUA01051582.1|:26-358

QVAKVTQRQC VLANRGVPQG GSLSGALCEL YYSAMLRSVD DFLYVTPSSS QANAFLTRMR

KGFPDFNCFI NKRKTQHNL- ---------- ---------- ---------- ----------

>gb|GETZ01034817.1|:1073-1984

QVAKVTQRQC VLANRGVPQG GSLSGALCEL YYSAMLRSVD DFLYVTPSSS QANAFLTRMR

KGFPDFNCFI NERKTQHNLR VAFRGIVMCS ESRQLQQVSL ARLPQVVLDP NIWRAGFMAG

>gi|1078083051|emb|HAES01102988.1|:C330-4

QIIRAGRNSR LLVTIGIAQG ATLSSELCEL YISAILRSVD DFLYITPSKE LAIYFMKVMK

EGFPDFNCFI NTSKT----- ---------- ---------- ---------- ----------

>gi|1077778384|emb|HAEZ01063549.1|:C1381-548

QIIRVGKNSR LLVTTGISQG STLSSELYEL YVSAMLRSVD DFLYITPSRE LAIYFMKIMN

EGFPDFNCII NSSKTQHNIR IIFRGTIFCT ASKQLNQIAK ARMPALLYDP NIWRCALSVG

>gi|1077778383|emb|HAEZ01063550.1|:C1381-548

QIIRVGKNSR LLVTTGISQG STLSSELYEL YVSAMLRSVD DFLYITPSRE LAIYFMKIMN

EGFPDFNCII NSSKTQHNIR IIFRGTIFCT ASKQLNQIAK ARMPALLYDP NIWRCALSVG

>gi|1128346665|emb|HAFG01037722.1|:C536-195

QIVRVGKNSI LLVTKGIAQG STLSSELYEL YVSAMLRSID DFLYITPSRE LAIYFMKIMN

EGFPDFNCII NSSKTQHNI- ---------- ---------- ---------- ----------

>gi|1076134614|emb|HAEW01003421.1|:C898-236

---------- ---------- ---------- -----LRSID DFLYITPSRE LAIYFLKVMN

EGFPEFNCFI NGTKTQHNIR IIFRGIVFCT ATKQLNQITK ARLPLLMVDP NIWRCSLSVG

>gi|1075680064|emb|HAEX01051271.1|:C846-121

---------- -------AQG ATLSSELYEL YISALLRSVD DFLFITHSRE LATYFLKVMN

EGFPEFNCYI NDTKTLHNIR IIFRGIVFCT ATKQLNQITK ARLPPLMFDP NIWRCSLSVG

>lcl|JW968276.1

QVIKIDSNY- YLQVKGIPQG GCFSSALSGI YYGHLFRAAD DFLFVTTLED LAEKFLVKAE

EGFPDYGCQI NRSKTRTNLR LP-------- ---------- ---------- ----------

>lcl|Ppruinosus29004

QLFQIGVRQR FLIVKGIAQG GSLSSALCDL YYSAMMRSVD DFLYITSRKT EAKNFLCFME

QGIDEFNCQI NPAKTMHNLR IPYRGTVFCP ASNQLKQILK SRFTKLISHP NIWRMGLIIG

>lcl|Pdispar36827

QLFQIGVRQR FLIVKGIAQG GSLSSSLCDL YYSAMMRSVD DFLYITSRKT EAKNFLRFIK

QGIDEFNCHI NPTKTMHNLR IP-------- ---------- ---------- ----------

>lcl|Trathkei36625

QLFQIGVRQR FLIVKGIAQG GSLSSALCDL YYSAMMRSVD DFLYITSKKT EAKTFLSLME

QGIDEFKCYI NPTKTMHNLR VPYRGTMFCP ASNQLHQ--- ---------- ----------

>lcl|Adrepressum54905

QLFQIGVRQR FLIIKGIAQG GSLSSDLCDL YYSAMMRSVD DFLYITPMKT QAEEFLRFME

QGIDEFNCHI NPGKTMHNL- ---------- ---------- ---------- ----------

>gb|GEZX01076726.1|:C1522-629

QRVSAGKKRT FLVTKGIAQG GAMSIDLCDL YYCALVRCVD DFLFISSDKT SATKFLDISS

KGVPEFNCYI NQSKTLHNVR VPFCGITVCS LTRQLEQITL ARLKDLVIHP NIYRVGIMTG

>gb|GDUJ01029506.1|:1765-2382

KYVSAGNNRK YRVTKGIAQG SFLSSALCDM YYSSMMRSVD DFLFVSVEKT SAEKFLAIAS

KGVPEYNCYI NARKTLQNIR VPFCGITLCS HSRQLLLITN ARLRVEVFHP NIYRTGMSVG

>gb|GEHV01052971.1|:C633-13

QYISAGRNRK YLITKGIAQG GFLSSTLCEL YYTSMLHSVD DFLFVSPEKI SAKRFLDVAA

RGVPDYNCVI NQTKTLHNIR VTFCGVTFCS FTRQLQQIAK TRLSAEVIHP NLHRTGISVG

>gb|GERA01033500.1|:1-303

QFISAGHNRK YSVTKGIAQG GSISSVLCEL YYNAMLHSVD DFLLISPCKI SAKKFLSVSM

KGVNGFNCFI NPNKTYHNI- ---------- ---------- ---------- ----------

>gb|GEPZ01015406.1|:470-913

QYISAGLNRK YLVTKGIAQG GTISSDLCEL YYNAMLHSVD DFLFISPIKH YAERFLELTV

NGVDQFNCFI NPKKTLHNIR VPFCGITICS LSRQ------ ---------- ----------

>gb|GERB01081677.1|:4-447

QYISAGINRN YLVTKGIAQG GTISSDLCEL YYNAMLHSVD DFLFISPIKR SAERFLELTV

KGVDQFNCFI NPKKTLHNIR VPFCGITICS LSRQ------ ---------- ----------

>gb|GEQR01035916.1|:C445-2

QYISAGLNRK YLVTKGIAQG GTISSDLCEL YYNAMLHSVD DFLFISPIKR SAECFLELTV

KGVDQFNCFI NPKKTLHNIR VPFCGITICS LSRQ------ ---------- ----------

>gb|GEPW01019622.1|:C513-103

QYISAGLNRK YLVTKGIAQG GTISSDLCEL YYNAMLHSVD DFLFISPIKR SAERFLELTV

KGVDQFNCFI NPKKTLHNIR VPFCGITICS LSRQ------ ---------- ----------

>gb|GEQA01015905.1|:C960-28

QYISAGLNRK YLVTKGIAQG GTISSDLCEL YYNAMLHSVD DFIFISPIKR SAEHFLELTV

KGVDHFNCFI NPKKTLHNIR VPFCGITICS LSRQLQQVTL ARLPDTVIHP NVYRVGLMAG

>gb|GESQ01009450.1|:4-738

-----GLNRK YLVTKGIAQG GTISSDLCEL YYNAMLHSVD DFLFISPIKR SAEHFLELTV

KGVDQFNCFI NPKKTLHNIR VPFCGITICS LSRQLEQVTL ARLPDTVIHP NVYRVGLMAG

>gb|GEQP01042736.1|:C998-201

QYISAGINKK YLVTKGIAQG GTISSMLCEL YYNAMMHCVD DFLFISPNKL SAKRFLELSV

KGVDQFNCFI NPKKTIHNIR VPFCGITICS LSRQLKQVTL ARLPDTVIHP NVYRVGLMAG

>gb|GEQE01032802.1|:1380-2309

QYISAGINKK YLVTKGIAQG GTISSMLCEL YYNAMMHCVD DFLFISPNKL SAKRFLELSV

KGVDQFNCFI NPKKTIHNIR VPFCGITICS LSRQLKQVTL ARLPDTVIHP NVYRVGLMAG

>gb|GEQK01024232.1|:1209-2006

QYISAGINKK YLVTKGIAQG GTISSMLCEL YYDAMLHSVD DFLFISPNKL SAERFLERCV

KGVDQFNCFI NPKKTIHNIR VPFCGVTICS LSRQLKQVTL ARIPDTVIHP NVYRVGLMAG
