## Supplementary material S3 for "The crustacean *Armadillidium vulgare* (Latreille, 1804) (Isopoda: Oniscoidea), a new promising model for the study of cellular senescence"

### Supplementary material 3

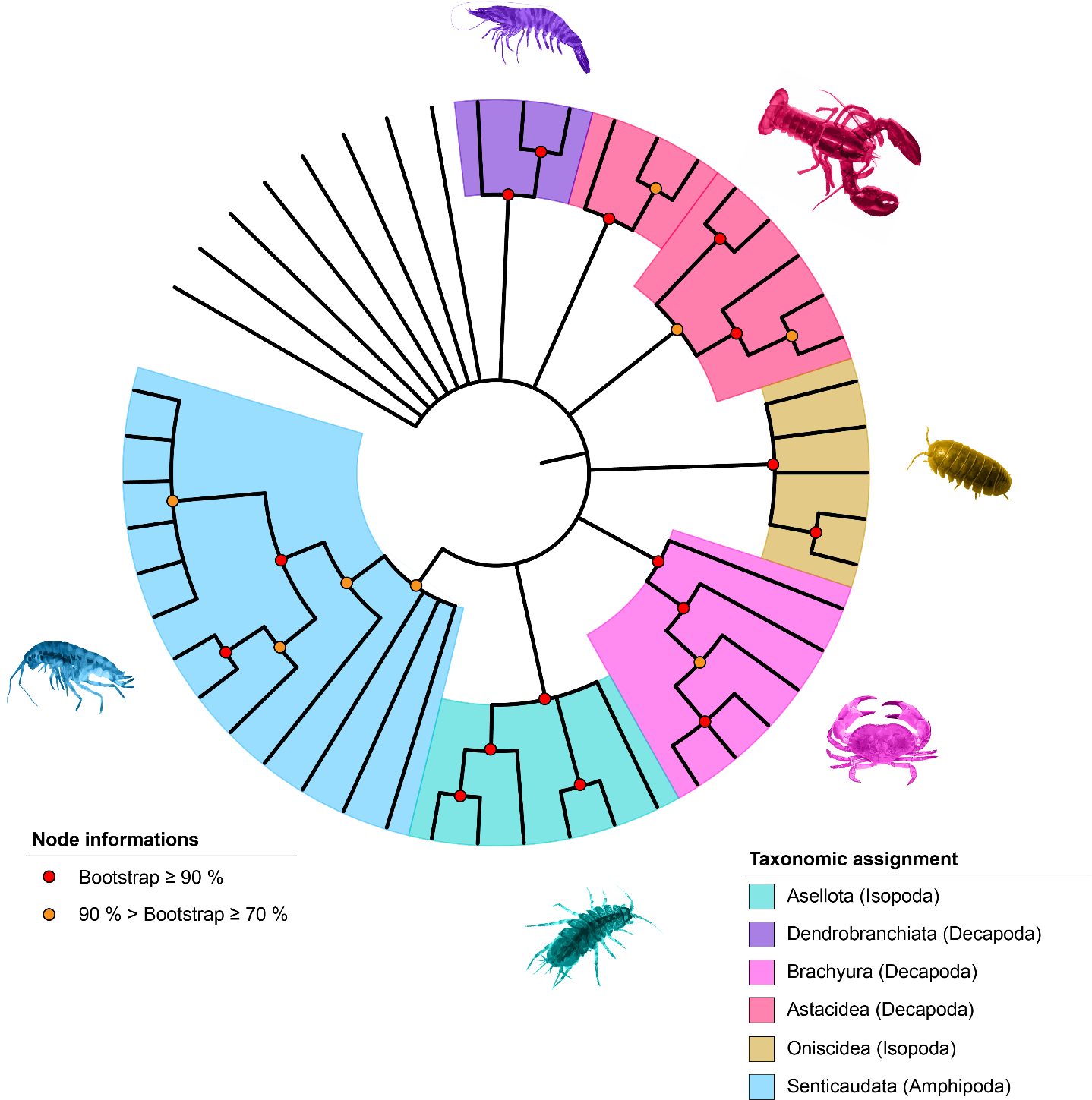

Phylogenetic analysis of the TERT homologous sequences identified in crustaceans (49 sequences).The molecular phylogenetic tree was inferred by the Maximum Likelihood method, based on 120 amino acids (LG + G model).
