## Supplementary material S4 for "The crustacean *Armadillidium vulgare* (Latreille, 1804) (Isopoda: Oniscoidea), a new promising model for the study of cellular senescence"

### Supplementary material 4

(((((((((((((AV_10400-RA:0.00000000,|Adrepressum54905:0.01337909):0.47018116[0.9930],|Ppruinosus29004:0.07662106):0.00000000[0.3970],|Trathkei36625:0.09808897):0.01135438[0.3640],|Pdispar36827:0.02943920):0.56137545[0.9520],('gb|GFFH01034216':0.75400494,|IABX01122151.1:0.56184732):0.00000000[0.4140]):0.00000000[0.1460],('gi|1078083051|emb|HAES01102988.1|330-4':0.04569749,(('gi|1076134614|emb|HAEW01003421.1|898-236':0.04216313,'gi|1075680064|emb|HAEX01051271.1|846-121':0.07691375):0.08746157[0.6860],('gi|1128346665|emb|HAFG01037722.1|536-195':0.04401228,('gi|1077778384|emb|HAEZ01063549.1|1381-548':0.00000000,'gi|1077778383|emb|HAEZ01063550.1|1381-548':0.00000000):0.02438274[0.9850]):0.09098611[0.9040]):0.06016991[0.4550]):0.48227972[0.9800]):0.10620582[0.0550],(('gb|GEZX01076726.1|1522-629':0.33353039,'gb|GDUJ01029506':0.36494830):0.00000000[0.3980],('gb|GEHV01052971.1|633-13':0.31748854,('gb|GERA01033500':0.25821294,((('gb|GEQP01042736.1|998-201':0.00000000,'gb|GEQE01032802':0.00000000):0.02337012[0.9910],'gb|GEQK01024232':0.04835881):0.03737848[0.8950],('gb|GEPZ01015406':0.03647857,('gb|GERB01081677':0.02355540,('gb|GEQA01015905.1|960-28':0.02970998,('gb|GEPW01019622.1|513-103':0.00000000,('gb|GEQR01035916.1|445-2':0.01091071,'gb|GESQ01009450':0.00682843):0.00966212[0.1790]):0.00000000[0.2510]):0.00000000[0.4080]):0.00000000[0.4380]):0.05161382[0.8950]):0.06566494[0.9000]):0.18838173[0.8340]):0.05191707[0.4330]):0.28914388[0.8860]):0.10032296[0.0530],(((('gb|GEUA01051582':0.01294608,'gb|GETZ01034817':0.00000000):0.00000000[0.9070],'gb|GFRT01019308.1|999-88':0.06067990):0.42268889[0.9910],'gb|GETT01116158.1|1474-671':0.41622352):0.09180984[0.3540],(('gi|1066151538|emb|HACK01015471':0.00000000,'gi|1068206298|emb|HACB02024865.1|917-87':0.00000000):0.27539321[1.0000],('gb|GBEI01054168.1|570-235':0.04114171,('gi|703876552|gb|GBEV01020382.1|1209-304':0.00000000,'gi|742973106|gb|GARH01006483.1|786-1':0.00000000):0.03716835[0.7920]):0.12845911[0.9830]):0.43823839[0.8390]):0.06710883[0.2140]):0.05499721[0.1760],((('gb|GFDA01078771.1|429-103':0.00000000,'gb|GFUC01026601':0.00000000):0.00000000[0.7160],'gb|GEBG01023904.1|1377-466':0.00000000):0.68463955[1.0000],('gb|GEFB01013694.1|752-144':0.51505596,('gb|GGFD01023608':0.19441513,('gb|GFFJ01110714':0.12092079,('gi|936323717|gb|GDRN01098857':0.00000000,('gi|936323711|gb|GDRN01098861':0.00000000,'gi|936323715|gb|GDRN01098858':0.00000000):0.00000000[0.3120]):0.22990682[1.0000]):0.09190091[0.8570]):0.32336901[0.9620]):0.73409968[0.9520]):0.20724875[0.2840]):0.42812791[0.3830],'gb|GEHY01011681.1|2384-2022':1.17158885):0.00000000[0.2250],('gb|GFUD01030199':1.11691760,'gi|595481066|gb|GARW01005962.1|1387-1037':0.55353587):0.49296988[0.3340]):0.00000000[0.3320],|JW968276.1:0.82130669):2.65542715[0.0000],NP_001035796.1:0.34338825)OROOT;
